# Concept design of a bionic grasp to capture the hand motion

**DOI:** 10.1101/2021.03.03.433703

**Authors:** Monirul Islam, Xinrui Li, Mahmoud Chizari

## Abstract

This study has focused on designing a device allowing the user to perform hand exercise motions, in relation with previous methods with good effects in stroke rehabilitation. The project is mainly aimed at designing a low cost, easy to handle and easy to manufacture smart glove to assist users with finger flexion and extensions, which can at the same time transfer data to an application. The methodology used here includes research, design, simulation testing and finalised design. It started with research and investigations on the needs of the users and previous efficient stroke rehabilitation methods. This has been followed with 4 concept designs, which has been tested and evaluated in the next stage for possible optimizations using a computer modelling technique. At the final step, the virtual prototype and product design specification (PDS) of the finalise concept for the interactive device has been illustrated.

## 1. Target application; stroke patients with physical challenges in the hand

Stroke, as one of the most life threating conditions that affects the human body, takes place when the fully oxygenated blood supply entering the brain is restricted and as a result, the brain cells will begin to die after a few minutes. [1] More than 100,000 strokes happen every year in the UK. In 2016, almost 38,000 people died from the causes of a stroke. [2] Studies conducted by the Stroke Association, 80% of survivors have a physical disability. Attending rehabilitation is the most important step in the recovery process in order to regain functionalities and strengthen muscles of the patients. It will normally comprise of: ‘physical activities, technology-assisted physically activities, cognitive and emotional activities. [3] This project was conducted after witnessing how many strokes occur every year, and the severe effects it puts onto the surviving patients. It will be mainly targeted at surviving stroke patients with physical challenges in the hand and is expected to help the patient with exercises for the hand to regaining the motor function in the brain for the duration of the rehabilitation programme.

### 1.1 The anatomy and mechanics of the hand

A human hand contains a system of muscles and tendons for complex mechanics in order to perform any functional movement, as in Figure 1.1 (a) and (b). The muscles can be divided into two groups: extrinsic, which are the long flexors on the underside of the forearm and puts together the tendons to the phalanges of the finger, and intrinsic. The deeper flexor is joined to the distal phalanx, while the superficial flexor is attached to the middle phalanx. [4] The human hand comprises of 27 bones. 8 of them are the carpal bones, 5 of them are the metacarpal bones and finally 14 bones in each finger. Each finger is made up of three bones with three joints which allows each finger to freely bend and extend in one way. The thumb is made up of two bones and only has two joints. [5]

**Figure 1.1:**
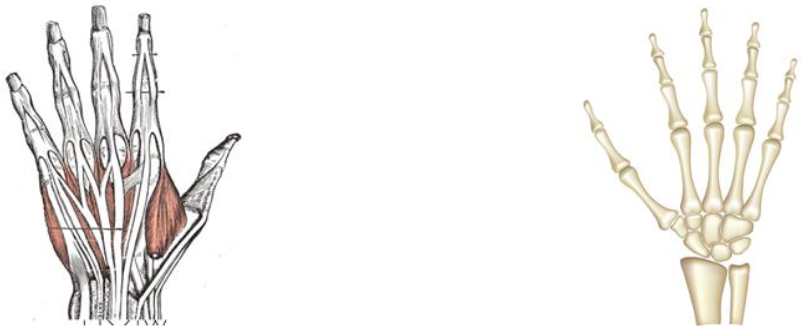
(a)Extensor compartments of wrist [4]; (b) Human hand Bone structure [5]

### 1.2 Rehabilitation background and current rehabilitation solutions

Occupational therapy assists surviving stroke patients to perform repetitive tasks and relearn their motor functions that were removed when the brain was damaged to help build strength and skill in daily life. These tasks can be little as washing the dishes or folding the laundry, but the more extensive approach can benefit the patient such as closing and opening their hand or moving objects. The level of intensity is determined by the patient’s functionality and the doctor’s assessment. [7] During the rehabilitation programme, the patient plan may include physical activities. Motor skill exercises can progress with muscle coordination and strength. Constraint induced therapy is when the healthy limb is contained, and the patient exercise the unhealthy limb in motion to help recover its motor function. Mobility training uses mobility supports such as walkers, cranes and wheelchairs. This helps balance out the bodyweight while the motor function in the leg helps the patient to walk. Finally, the range of motion therapy are specific movements that can comfort muscle tension and help reclaim its motion function. [3] Occupational therapy works with all ages at home or a clinic and those who have difficulties with the day-to-day life due to a physical or learning disability. [8] Home rehabilitation is very beneficial for mental and functional performance. The earlier rehabilitation starts, then the quicker the patient will be able to regain the motor skills through repetition of the exercises and mental motivation. [3] There are existing solutions to help patients regain their motor functions to the dominant state, such as a wearable glove to exercise their fingers to gain muscle strengths and coordination. Another solution is a wearable device, where the user is mainly using it for measurements (muscles movement) and see the progression throughout their rehabilitation process. The most beneficial for the patient to use is the wearable interactive product, as this will ensure the user actively exercise their muscles at different tensions. For example, SaeboGlove, designed by Saebo, is a functional object which allows neurological and orthopedic patients to help extension of their finger and thumb. [9] It is a soft glove with rubber tensioner to help with expansion, on which the finger and thumb extension are designed with interphalangeal tensioners for each joint. A silicon liner at each fingertip to improve traction during grasping. A unique wrist design to support the wrist. The palm is exposed to increased breathability. [10]

### 1.3 Reviews of existing sources

After discussing each individual source on the aim, main consideration, relevance to this project and limitation, as in Table 1.1, the project design will mainly focus on designing a mechanically-driven, lightweight, portable and adjustable glove.

**Table 1.1:**
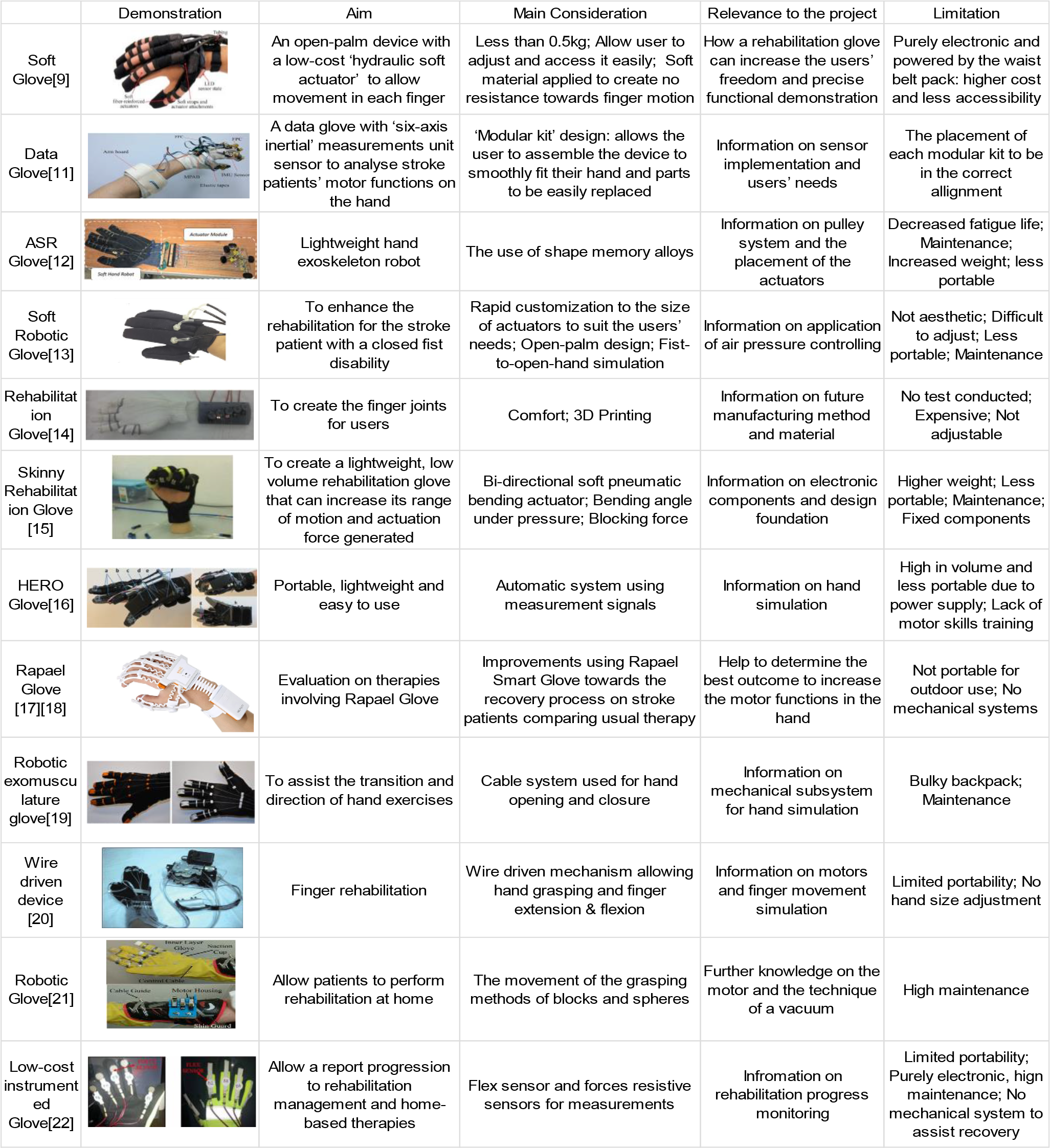
Reviews of 12 relevant sources

|  | Demonstration | Aim | Main Consideration | Relevance to the project | Limitation |
| --- | --- | --- | --- | --- | --- |
| Soft Glove[9] |  | An open-palm device with a low-cost 'hydraulic soft actuator' to allow movement in each finger | Less than 0.5kg; Allow user to adjust and access it easily; Soft material applied to create no resistance towards finger motion | How a rehabilitation glove can increase the users' freedom and precise functional demonstration | Purely electronic and powered by the waist belt pack; higher cost and less accessibility |
| Data Glove[11] |  | A data glove with 'six-axis inertial' measurements unit sensor to analyse stroke patients' motor functions on the hand | 'Modular kit' design: allows the user to assemble the device to smoothly fit their hand and parts to be easily replaced | Information on sensor implementation and users' needs | The placement of each modular kit to be in the correct alignment |
| ASR Glove[12] |  | Lightweight hand exoskeleton robot | The use of shape memory alloys | Information on pulley system and the placement of the actuators | Decreased fatigue life; Maintenance; Increased weight; less portable |
| Soft Robotic Glove[13] |  | To enhance the rehabilitation for the stroke patient with a closed fist disability | Rapid customization to the size of actuators to suit the users' needs; Open-palm design; Fist-to-open-hand simulation | Information on application of air pressure controlling | Not aesthetic; Difficult to adjust; Less portable; Maintenance |
| Rehabilitation Glove[14] |  | To create the finger joints for users | Comfort; 3D Printing | Information on future manufacturing method and material | No test conducted; Expensive; Not adjustable |
| Skinny Rehabilitation Glove [15] |  | To create a lightweight, low volume rehabilitation glove that can increase its range of motion and actuation force generated | Bi-directional soft pneumatic bending actuator; Bending angle under pressure; Blocking force | Information on electronic components and design foundation | Higher weight; Less portable; Maintenance; Fixed components |
| HERO Glove[16] |  | Portable, lightweight and easy to use | Automatic system using measurement signals | Information on hand simulation | High in volume and less portable due to power supply; Lack of motor skills training |
| Rapael Glove [17][18] |  | Evaluation on therapies involving Rapael Glove | Improvements using Rapael Smart Glove towards the recovery process on stroke patients comparing usual therapy | Help to determine the best outcome to increase the motor functions in the hand | Not portable for outdoor use; No mechanical systems |
| Robotic exomusculature glove[19] |  | To assist the transition and direction of hand exercises | Cable system used for hand opening and closure | Information on mechanical subsystem for hand simulation | Bulky backpack; Maintenance |
| Wire driven device [20] |  | Finger rehabilitation | Wire driven mechanism allowing hand grasping and finger extension & flexion | Information on motors and finger movement simulation | Limited portability; No hand size adjustment |
| Robotic Glove[21] |  | Allow patients to perform rehabilitation at home | The movement of the grasping methods of blocks and spheres | Further knowledge on the motor and the technique of a vacuum | High maintenance |
| Low-cost instrumented Glove[22] |  | Allow a report progression to rehabilitation management and home-based therapies | Flex sensor and forces resistive sensors for measurements | Information on rehabilitation progress monitoring | Limited portability; Purely electronic, high maintenance; No mechanical system to assist recovery |

**Table 1.2:** Product Design Specification

| Specification | Requirements |
| --- | --- |
| Dimensions (cm) | 18x20.3x0.5 – 25x26x2 |
| Weight (kg) | Maximum 0.5 |
| Volume (cm<sup>3</sup>) | 182.7 – 1300 |
| Ease of assembly | Must be easy to assemble and disassemble |
| Number of Components | To minimize complexity and weight no more than 40 components |
| External tools needed | Dependent on the function |
| Ease of Use | Very easy to operate the device, must be adjustable |
| Performance | Allows the user to flex and extend each finger while exercising the hand to regain motor function |
| Strength of design | Design must be strong enough so no damages can be inflicted to the device or users' hand |
| Aesthetics | Not too many loose wirings and must look in fashionable |
| Meets Aims Functions | The device allows the user to exercise their fingers and hand |
| Operate in different temperatures (°c) | -55 to 110 |
| Users Safety | Chamfer edges to 0.01 mm– 0.2mm |
| Material | ABS, Elastic string |
| Environmental safety | ABS can be recycled if you want to re-heat the material to use it again<br>Elastic string are organic cotton and natural rubber |
| Ease of manufacture | 3D printed or laser cutter |
| Cost of manufacture | ABS per cm<sup>3</sup>- £0.20<br>Elastic string for 2mm x 5m = £2 |
| Product cost | £15 - £50 |
| Maintenance | The device must be easy to clean and no wear and tear |
| Replaceable parts | Parts can be replaced if damaged |
| Lifespan | 5 year+ depending on usage |

### 1.4 Product Design Specification (PDS)

A general PDS was produced to establish important requirements and guidelines for each design.

## 2 Concept Design

### 2.1 Initial Concept Designs

To address the proposed research target, several design ideas were thought and sketched. Four of the completed sketches have been illustrated in Figure 2.1.

**Figure 2.1:**
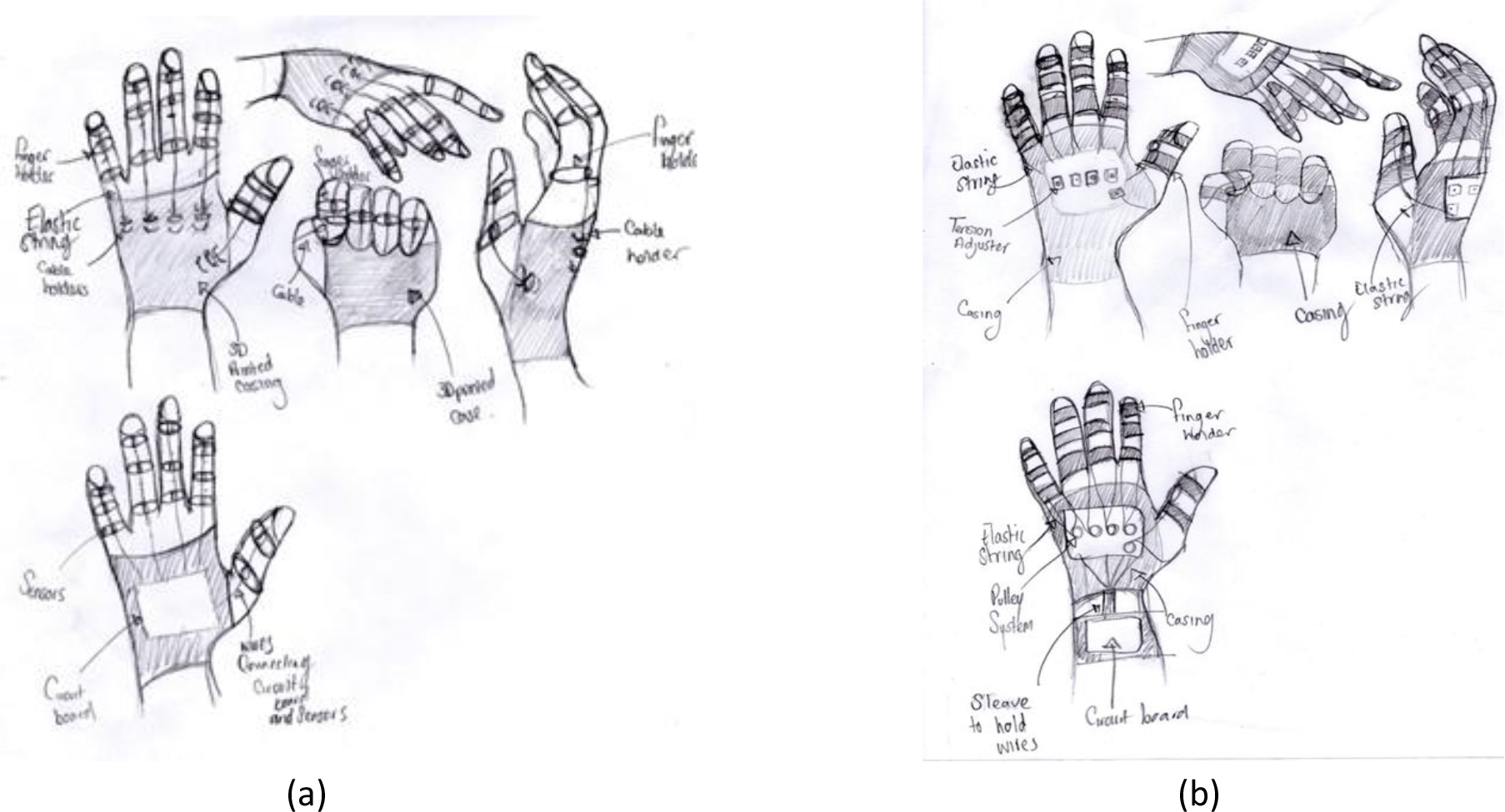
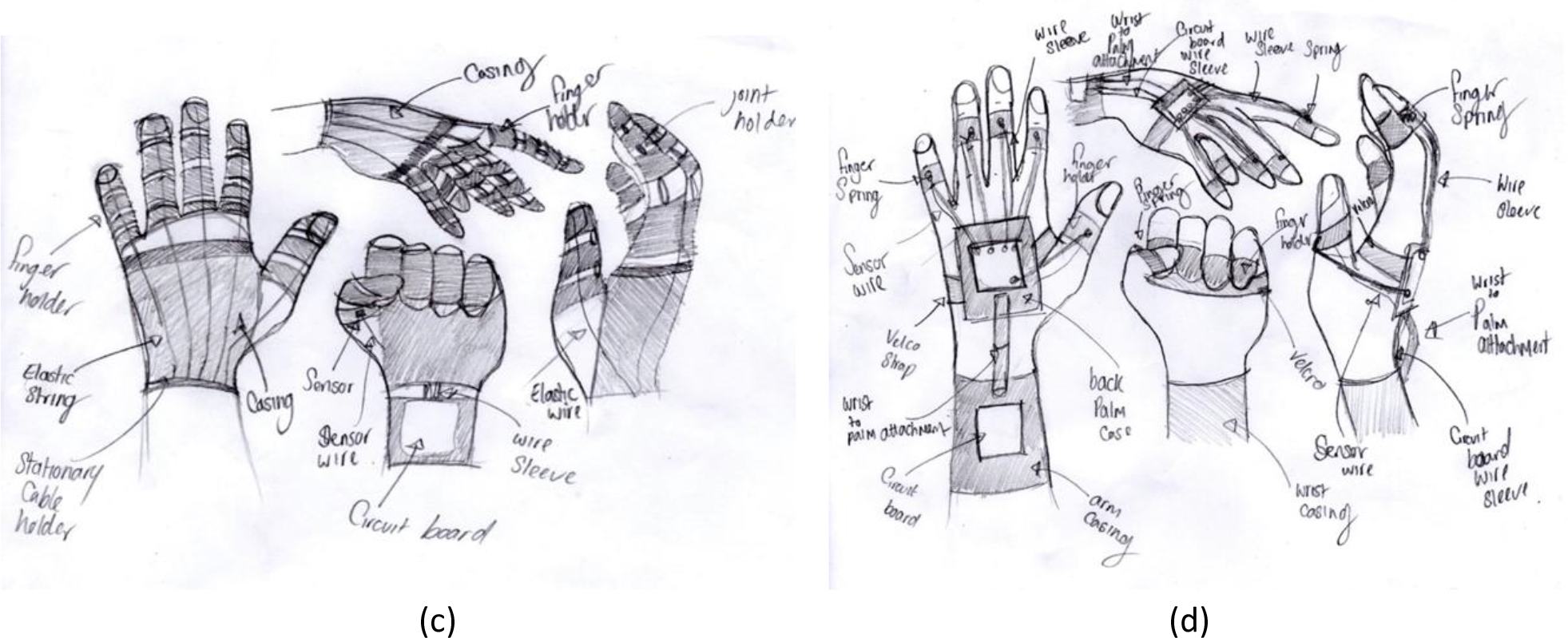
(a)Concept Design 1; (b)Concept Design 2; (c) Concept Design 3; (d) Concept Design 4

### 2.2 Evaluation of Concept 1

This design in Figure 2(a) includes a 3D printed ABS material casing for the hand, with three different tensioner level slots for each finger. The elastic string is to be placed to the corresponding level slot depending on the amount of tension required. The furthest slot from the finger would create a higher tension in the elastic string and generate more resistance when extending and flexing each finger. Slots near the corresponding finger would form minimal tension in the elastic string and create less resistance in the movement of each finger. This would allow the user to freely choose an exercise resistance level that has been advised by the rehabilitation officer/doctor. Each finger has a three ABS circular finger rings to hold the glove in place and the elastic string to pass through each ring to simulate a ‘tendon’ through each finger. Whereas, the thumb only contains two thumb rings. A circuit board would be placed on the surface of the hand inside the casing to allow to send data on the performance of the user motion to the selected app on any device its installed on. This design would be lack of adjustability and comfort due to 3D printed specific-to-user-size casing fitting the whole hand. The circuit board will require a power source, external or internal, which may increase the weight and require more energy to move to perform any motion.

### 2.3 Evaluation of Concept 2

This concept design in Figure 2(b) includes a 3D printed ABS casing for the hand, with five different tensioner adjusters on the surface of the hand casing. Two elastic stings are placed on the sides of each finger ring and coiled around the pulley system within the case. This will allow the user to change the tension of each wiring to the needed resistance. This gives the user more of a flexible option to choose from. Each finger has three circular finger rings to hold the glove in place and the elastic string passes through each ring to simulate a tendon through each finger. Whereas, the thumb only contains two thumb rings. A circuit board would be placed on the arm underneath the wrist inside the casing, this will allow to send data on the performance of the user motion to the selected app on any device its installed on. For this design, it would be also be very difficult for adjustability as the casing is 3D printed specific to the users’ size. Also, the circuit board will require a power source. This may increase the weight of the glove. As the circuit board is underneath the wrist, this may restrict some wrist movement because of the ABS material is not as flexible. To change the tension of each string, this would require an external tension adjuster to be slotted through the hole of the mechanical system. The elastic string is placed on the sides of each finger, this may require more force on the strings to be able to extend and flex the finger. This is because the wiring is not placed at the centre of mass point to create the maximum performance.

### 2.4 Evaluation of Concept 3

This concept design in Figure 2(c) involves a 3D printed ABS material casing for the hand, with a fixed-point elastic sting holder at the bottom of the case. This will allow the user to have one tension to be comfortable with and show improvements based on one tension throughout the rehabilitation process. Each finger has four ABS circular finger rings to hold the glove in place. An elastic string passes through each ring to simulate a ‘tendon’ through each finger. Whereas, the thumb only contains three ABS thumb ring. A circuit board would be placed on the back side of the arm underneath the wrist inside the casing, this will allow to send data on the performance of the user motion to the selected app on any device its installed on. For this design, it would also be very difficult for adjustability for the user, as the casing is 3D printed specific to the users’ size. Also, the circuit board will require a power source. This may increase the weight of the glove. As the circuit board is underneath the wrist, this may restrict some wrist movement because of the ABS material is not as flexible. Due to only one tension available on the glove, this will make progression slower. Once the user muscle is comfortable to the resistance, the use of this glove is not required.

### 2.5 Evaluation of Concept 4

This concept design in Figure 2(d) involves a 3D printed ABS material casing for the only the surface of the hand which is connected by a Velcro strap for adjustability. There are five fixed tensioners on the surface of the hand casing. A wire sleeve that is attached to the casing and the finger strap which contains the elastic stings and sensor wiring. Each finger has one circular ABS finger rings to hold the glove in place and the elastic string to pass through each ring to simulate a tendon through each finger. The sensor wiring is also connected to the circuit board. The circuit board would be placed on the arm underneath the wrist inside the casing, this will allow to send data on the performance of the user motion to the selected app on any device its installed on. The circuit board will require a power source, this may increase the weight of the glove and make it difficult to perform exercising motions. Due to only having one tension available on the glove, this will make progression slower and once the user muscle is comfortable to the resistance, the performance of the device will decline.

### 2.6 Concept selection

Before any concept design was rated, the product design specification features were weighted for its importance in the project. With a weight of 1 results the feature is not as important in the project and could be continued without compared to a weight of 5 on which the feature is an essential part of the project. All the concepts were rated against the general product design specification as shown in Table 1.4. A rating from 1-5, with 1 being poor at meeting the condition of the PDS and 5 being excellent in meeting the requirement. As calculated in Table 2.1(b), concept 4 scored the highest weighted rating of a total of 320. However, even though concept 4 scored the highest, features from other concept has been taken into consideration for development. For example, tension adjuster for the elastic string from concept 2 will significantly improve the rate of the rehabilitation process. This will allow the user to change the tension of each string to the required resistance. This will give the user more of a flexible option to choose from. In majority of the concept designs a circuit board was placed to allow the device to send data on the performance of the user motion to the selected app. This feature was considered, however the limitation of a power source which can limit the weight and portability of the glove was a factor in the decision of the development. As the budget of the project was very low, purchasing a full coded application would exceed the budget. Also, the coding of the application would be extremely challenging as the knowledge is very minimal. Hence, the concept idea of a device to send data based on performance has been removed. Therefore, concept 4 would be developed further for the base final design with exceeded rating features from other concept designs.

**Table 2.1:** (a) Weight and rating standards, (b) Concept Selection Chart

|  |  | Concept 1 |  | Concept 2 |  | Concept 3 |  | Concept 4 |  |
| --- | --- | --- | --- | --- | --- | --- | --- | --- | --- |
| Features | Weight | Rating | Weighted score | Rating | Weighted score | Rating | Weighted score | Rating | Weighted score |
| Dimensions | 3 | 5 | 15 | 5 | 15 | 5 | 15 | 5 | 15 |
| Weight | 5 | 5 | 25 | 5 | 25 | 5 | 25 | 5 | 25 |
| Volume | 3 | 4 | 12 | 3 | 9 | 5 | 15 | 4 | 12 |
| Ease of assembly | 3 | 5 | 15 | 4 | 12 | 5 | 15 | 4 | 12 |
| Number of Components | 2 | 5 | 10 | 4 | 8 | 5 | 10 | 4 | 8 |
| No External tools needed | 1 | 5 | 5 | 4 | 4 | 5 | 5 | 4 | 4 |
| Ease of Use | 4 | 5 | 20 | 4 | 16 | 5 | 20 | 4 | 16 |
| Ease to store | 2 | 5 | 10 | 5 | 10 | 5 | 10 | 5 | 10 |
| Performance | 5 | 3 | 15 | 5 | 25 | 2 | 10 | 5 | 25 |
| Strong design | 3 | 4 | 12 | 5 | 15 | 4 | 12 | 5 | 15 |
| Aesthetics | 1 | 3 | 3 | 5 | 5 | 3 | 3 | 5 | 5 |
| Meets Aims Functions | 5 | 4 | 20 | 5 | 25 | 3 | 15 | 5 | 25 |
| Operate in different temperatures | 4 | 5 | 20 | 5 | 20 | 5 | 20 | 5 | 20 |
| Users Safety | 4 | 5 | 20 | 5 | 20 | 5 | 20 | 5 | 20 |
| Material | 5 | 5 | 25 | 5 | 25 | 5 | 25 | 5 | 25 |
| Environmental safety | 4 | 5 | 20 | 5 | 20 | 5 | 20 | 5 | 20 |
| Ease of manufacture | 2 | 5 | 10 | 4 | 8 | 5 | 10 | 4 | 8 |
| Cost of manufacture | 3 | 5 | 15 | 4 | 12 | 5 | 15 | 4 | 12 |
| Product cost | 2 | 5 | 10 | 4 | 8 | 5 | 10 | 4 | 8 |
| Maintenance | 3 | 3 | 9 | 3 | 9 | 3 | 9 | 5 | 15 |
| Replaceable parts | 1 | 2 | 2 | 4 | 4 | 1 | 1 | 5 | 5 |
| Lifespan | 3 | 3 | 9 | 3 | 9 | 3 | 9 | 5 | 15 |
| Sum of specification values |  |  | 302 |  | 304 |  | 294 |  | 320 |

| Weight |  | Rating |  |
| --- | --- | --- | --- |
| Very Important | 5 | Excellent | 5 |
| Fairly Important | 4 | Good | 4 |
| Moderately Important | 3 | Fair | 3 |
| Slightly Important | 2 | Sufficient | 2 |
| Not Important | 1 | Poor | 1 |

## 3 Final Design

### 3.1 Design Version 1: Evaluation

The Final design V1, showed in Figure 3.1 (a) and (b), involves a 3D printed ABS material case placed above a hand brace which is connected by a Velcro strap for adjustability. The elastic stings are placed inside of a plastic tubing attached to the casing and the ABS finger ring. The elastic wiring is coiled around the bearing pulley system within the case. This will allow the user to change the tension of each string to the needed resistance for motion. This gives the user a more flexible option to choose from during their rehabilitation process. Each finger has one circular ABS finger rings which can be easily slotted in by applying force to the designated area. Each ring will hold the glove in place with the plastic tubing. The elastic string will be placed inside the plastic tubing to simulate a tendon through each finger. The plastic tubing does not have much motion of freedom, for example if the user does a 90° motion, the tubes will not be able to fully bend. This will result in touching the users’ skin and create more friction with the elastic string.

**Figure 3.1:**
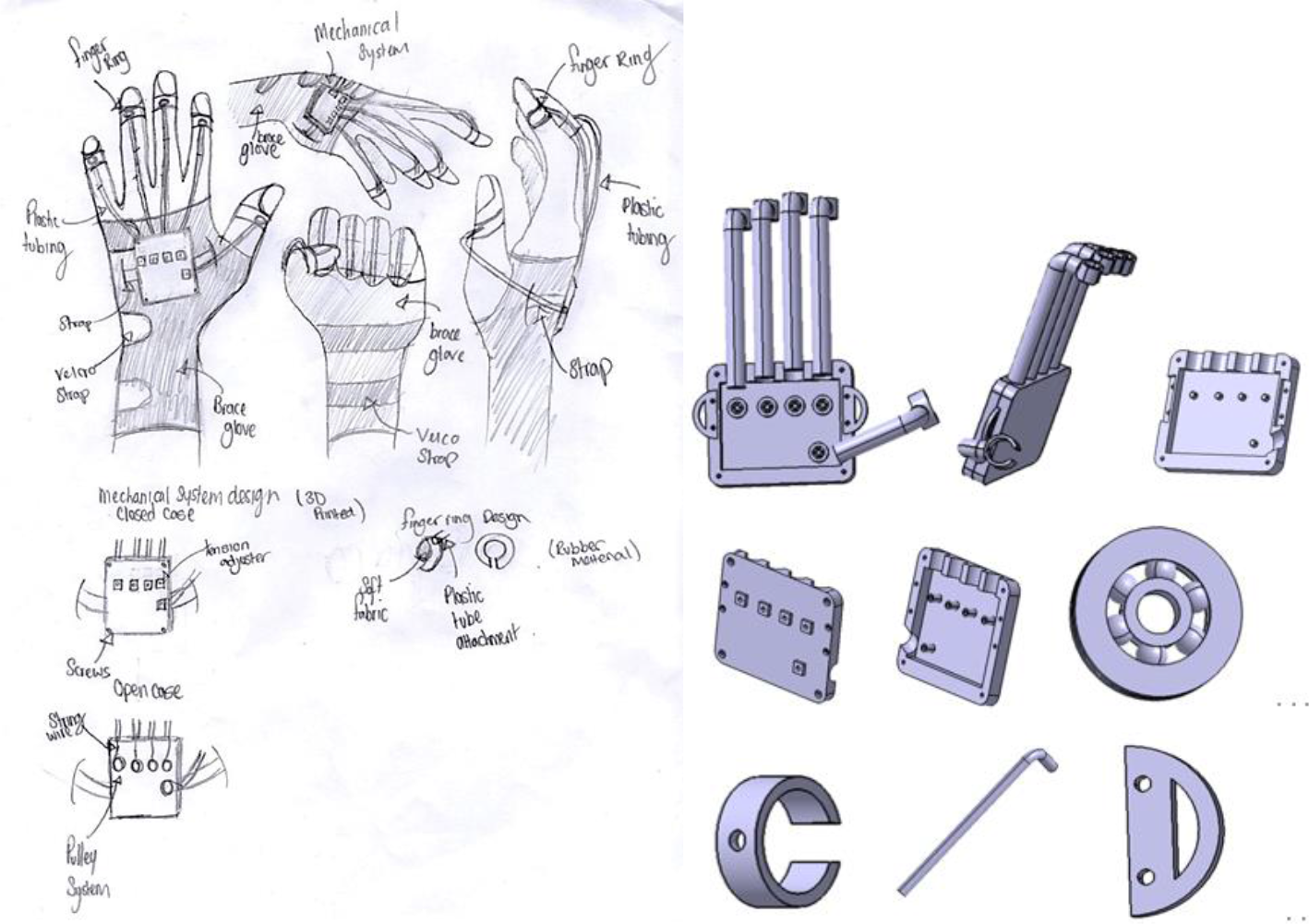
(a) V1 sketch; (b) V1 by Catia 3D: assembly, bottom case, bearing pully, finger ring, plastic tube and Velcro strap holder

The final design assembly consists of all the individual components excluding the hand brace and elastic string: Bottom casing, Top casing, 5 bearing pulleys, 4 finger rings, 4 plastic tubing, a thumb ring, a thumb plastic tube and 2 Velcro strap holders. Each plastic tubing will be slotted down depending on the users’ size information. Each elastic string will be coiled around to their corresponding bearing pulley and travel down to the end of each finger ring where a knot will be tied to secured in place. The tension can be changed of each elastic string by placing a circular key through the hole of the top case and slotted into the bearing pulley. Turning the key clockwise will make the elastic string will contract and coil around. This will create more resistance when extending and flexing the fingers. This will improve the rate of improve the motor function skill during the rehabilitation process, however it is advised not to force the resistance on a regular basis, as it may create more tears in the muscle. Only a trained therapist with experience should change the tensions.

From visual analysis, the plastic tubing does not have much motion of freedom and will not be able to perform a 90° motion, As the tubes bends, this will damage the tube overtime and also the finger joints will touch the tubing and create some friction on the elastic string or make the user feel uncomfortable while exercising.

### 3.2 Design Version 1: Further development

To improve the design for maximum performance, the plastic tubing must be changed to allow no resistance on the elastic string and allow freedom of motion for all angles.

### 3.3 Design Version 2: Evaluation

The following design in Figure 3.2 (a) and (b) has been 3D modelled on Catia V5, the individual components from the final design are still being used in this model however with some adjustments made to the plastic tubes which are shown excluding the hand brace and elastic wiring. The optimised design includes a 3D printed ABS material case which is placed above a hand brace and connected by a Velcro strap to allow adjustability. The elastic stings are placed through several plastic tubing acting as tendons in the finger attached to the casing and several finger rings for stability. The plastic ‘tendons’ are connected to each other plastic tube using a ball shaft. This will allow multiple degree of motion for extension and flexion with the fingers. As in the previous designs, the elastic string is coiled around the bearing pulley system within the case. Each finger has three circular finger rings which can be easily attached by applying force to place it in. Whereas the thumb contains two rings. Each ring will hold the glove in place with the plastic tubing. The elastic string will be placed inside the plastic tubing to simulate a tendon through each finger. The optimised design assembly consists of all the individual components from the final design excluding the hand brace and elastic string and plastic tubes: Bottom casing, Top casing, 5 bearing pulleys, 12 finger rings, 16 plastic tubing, 2 thumb ring, 3 thumb plastic tube and 2 Velcro strap holders. Each plastic tubing will be slotted down depending on the users’ size information. Each elastic string will be coiled around to their corresponding bearing pulley and travel down to the end of finger ring where a knot will be tied to secure in place. Each plastic tubing can be connected like Lego using the ball shaft as shown in Figure 3.2 (b). This will allow a different degree of motion for extension and flexion with the fingers. There is an attachment at the bottom of the plastic tubes to attach onto the finger ring, this is to allow no skin contact for the user to feel uncomfortable when performing exercises. As the same feature as the final design, the tension can be changed of each elastic string by placing a circular key through the hole of the top case and slotted into the bearing pulley. This will create more resistance when extending and flexing the fingers. This will significantly improve the rebuilding of the motor function skill during the rehabilitation process, however as stated in previous concept, it is advised not to force the resistance on a regular basis, as it may create more tears in the muscle. Only a trained therapist with experience should be allowed to change the tensions of the string. Table 3.1 demonstrates the PDS of this optimized version.

**Table 3_1:**
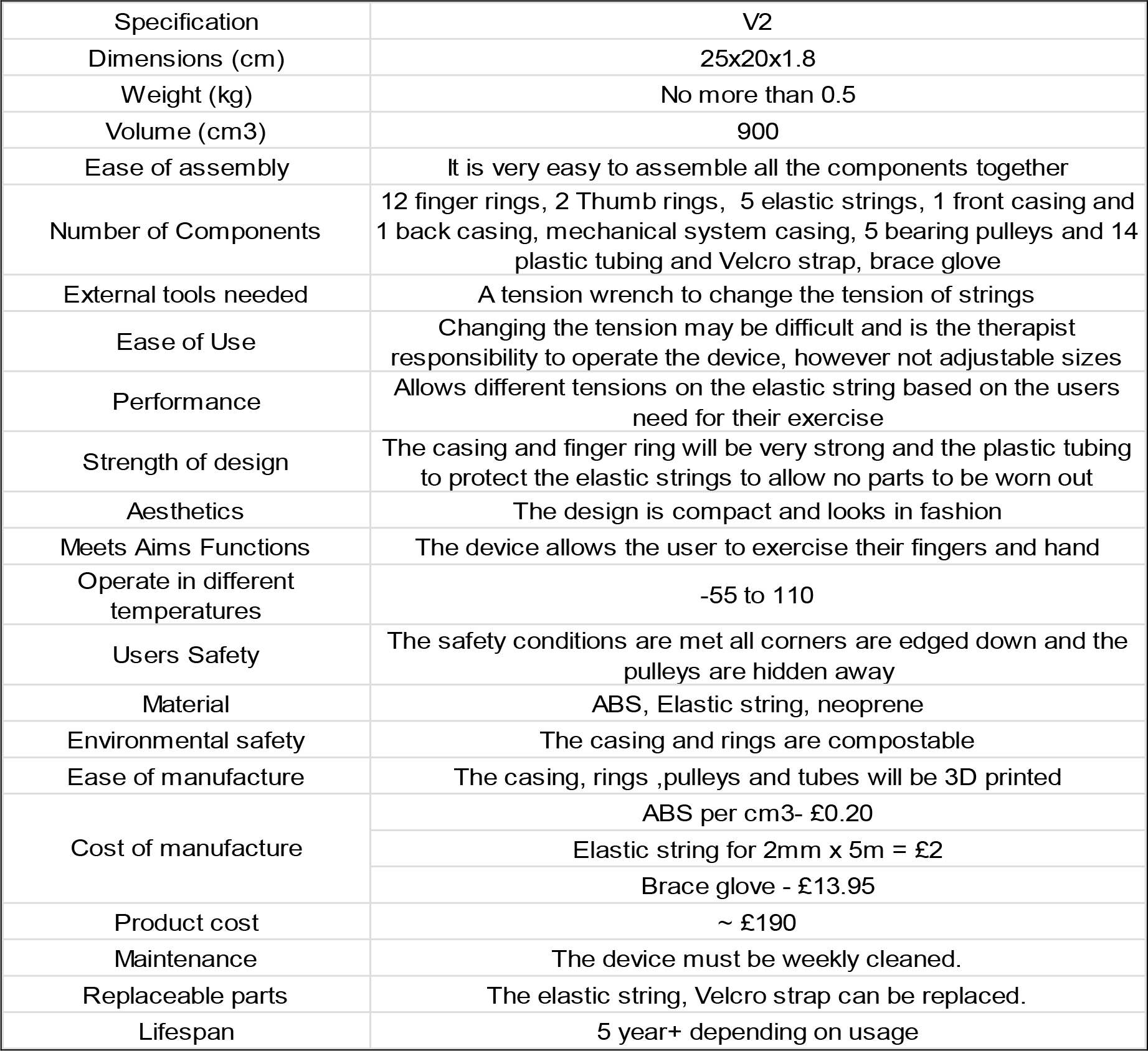
PDS of the final design version 2

**Figure 3.2:**
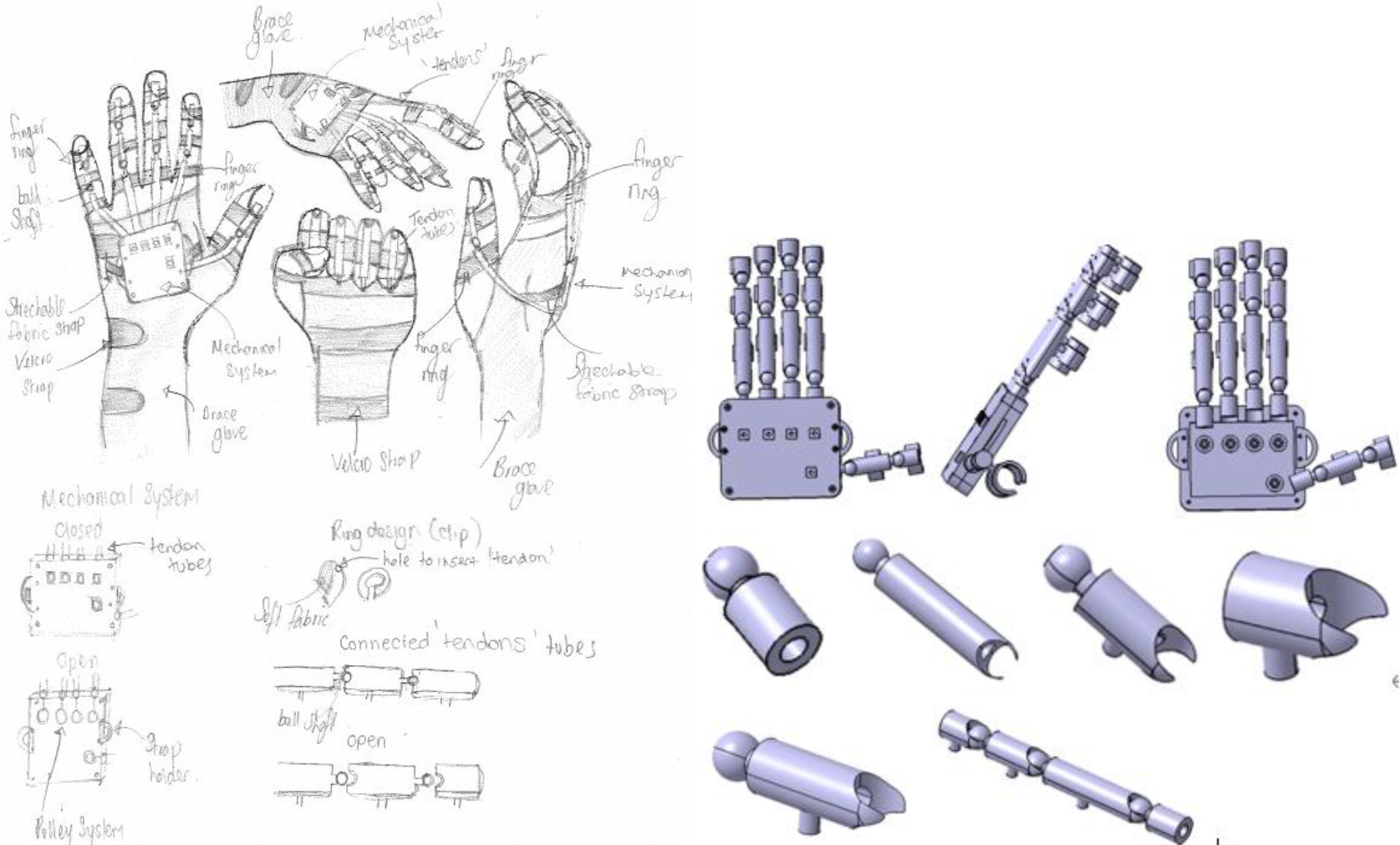
(a) V2 sketch; (b) V2 by Catia 3D: assembly, finger components

## 4 Conclusion

This research has been mainly targeted at surviving stroke patients with physical challenges in the hand and aimed to help the patient exercise to regain the motor function in the brain for the duration of the rehabilitation programme. Research on the rehabilitation background and reviews on current occupational rehabilitation solutions have been conducted for the following design process. 4 concept designs were presented and compared for the most suitable concept according to the PDS. Based on Concept 4, CAD model based on the selected concept design were created and analysed on Catia V5. Further developments for maximum performance have been decided and illustrated in the CAD model of the finalised design. It consists of a 3D printed ABS material case placed above a hand brace and connected by a Velcro strap, including bottom casing, top casing, 5 bearing pulleys, 12 finger rings, 16 plastic tubing, 2 thumb rings, 2 thumb plastic tubes and 2 Velcro strap holders. Different degrees of motion for finger extension and flexion, as well as no skin contact during use, have been realized. To compare with the original PDS, the finalised design has met most of the criteria and has a good agreement with authors’ provirus finding [23].

